# Structure of the T=13 capsid of infectious pancreatic necrosis virus (IPNV) - a salmonid birnavirus

**DOI:** 10.1101/2024.08.20.608766

**Authors:** Anna Munke, Amr Ahmed Abdelrahim Gamil, Aase B. Mikalsen, Han Wang, Øystein Evensen, Kenta Okamoto

## Abstract

Birnaviruses infect a broad range of vertebrate hosts, including fishes and birds, and cause substantial economic losses in the fishery and livestock industries. The infectious pancreatic necrosis virus (IPNV), an aquabirnavirus, specifically targets salmonids. While structures on T=1 subviral particles of the birnaviruses, including IPNV, have been studied, structural insights into the infectious T=13 particles have been limited to the infectious bursal disease virus (IBDV), an avibirnavirus. Determining the capsid structure of the T=13 particle of IPNV is crucial for advancing knowledge of its antigenicity, capsid assembly, and possible functional structures. Here, the capsid structure of the IPNV L5 strain has been determined at a resolution of 2.75 Å. The overall structure resembles the T=13 IBDV structure, with notable differences in the surface loops on the P domain of the VP2 capsid protein, essential for antigenicity and virulence. Additionally, previously undescribed structural features have been identified, including the C-terminal regions of the VP2 subunits within the pentagonal assembly unit at each 5-fold axis, which interlock with adjacent VP2 subunits. This interlocking, together with class-averaged projections of triangular and pentagonal units, suggests that the pentagonal unit formation could be important for correct T=13 particle assembly, preventing the formation of T=1 subviral particles. Furthermore, positively charged residues in obstructed capsid pores at each 5-fold axis are speculated to facilitate intraparticle genome synthesis of IPNV.

**Importance:** Aquabirnaviruses cause deadly infectious diseases in salmonid fish, posing significant challenges for both wild and farmed fish populations. The most prevalent aquabirnavirus worldwide is the infectious pancreatic necrosis virus, whose multifunctional capsid is critical to its infection, replication, and maturation. Previously, research has focused on the structure of the virus’s non-infectious subviral capsid. In this study, however, the first structure of the large, infectious, and functional form of the capsid has been determined. This new capsid structure reveals functional motifs that were previously unclear in the non-infectious capsid. These motifs are believed to be essential for the virus’s replication and particle assembly, making them promising targets for developing strategies to control virus proliferation.

## Introduction

*Birnaviridae* viruses are disease-causative agents to vertebrates, encompassing a few species in the genus *Aquabirnavirus,* infecting fishes, and genus *Avibirnavirus,* infecting birds. Recently, putative birnaviruses have also been identified in pigs (1). The infectious pancreatic necrosis virus (IPNV), a member of the *Aquabirnavirus*, can infect diverse salmonid species, leading to lethal outcomes characterized by pancreatic necrosis and catarrhal enteritis (2, 3). Because of its negative impact on farmed fish, control measures such as vaccination and genetic selection have been implemented to combat this virus (4, 5). Another important birnavirus pathogen, the infectious bursal disease virus (IBDV), infects chickens and turkeys, leading to severe immunosuppression (5, 6).

Birnaviruses have a bi-segmented dsRNA genome. Segment A encodes a large polyprotein consisting of pVP2-VP3-VP4, and, in some species, a short VP5 protein. Segment B encodes the RNA-dependent RNA polymerase (RdRp), VP1 (7, 8). The preVP2-VP3-VP4 polyprotein undergoes self-cleavage by the VP4 protease, generating preVP2 and VP3. During particle assembly, preVP2 is further processed to mature VP2 by cleaving its C-terminus region (8). The cleaved C-terminal fragments of VP2 are associated with the mature capsid (9). VP3s and VP1s form highly ordered interior ribonucleoprotein complexes (RNP) to package the segmented genomes inside the particle (10, 11), while VP2s assemble into a 60-70 nm T=13 icosahedral single layer capsid to protect the RNPs (12, 13). The precursor N- and C-terminal regions of VP2 and the C-terminal peptide of VP3 are critical for the formation of infectious large T=13 particles since expressing only the matured form of VP2 results in the formation of non-infectious small ∼23 nm T=1 subviral particles (13–18).

The crystal structures of T=1 subviral particles from both IBDV and IPNV have been determined, revealing three structural domains within VP2: the P, S, and B domains (13, 15, 16, 19). The S domain employs a typical jelly-roll fold of icosahedral viruses, which implies an evolutionary link between the +ssRNA nodaviruses and tetraviruses and the dsRNA birnaviruses and reoviruses (13, 20). The P domain, unique to birnaviruses, is considerably variable in amino acid sequence and forms surface protrusions (13, 16). Structural variations in the surface loops of the P domain between the IBDV and IPNV subviral particles likely play a role in regulating virulence and tropism (16). Certainly, mutations in the P domain often result in less virulent strains (21–24). The B domain, located inside the virus capsid, is believed to play a role in genome encapsulation through its arrangement of α-helices, as described in other ssDNA and +ssRNA viruses that employ the jelly-roll fold (20, 25–27). The T=1 icosahedron contains only one VP2 subunit in the asymmetric unit, and the structure of this VP2 has been resolved in a single conformation (16), limiting the understanding of the functional structures of the infectious T=13 capsid. This limitation also hinders the comprehension of possible mechanisms of T=13 particle assembly.

Another uncertain aspect of the birnavirus capsid structure is the surface pore at each 5-fold axis. This pore possibly has a role in facilitating the synthesis of virus transcripts inside the capsid by incorporating nucleoside triphosphates (NTPs) and releasing +ssRNA virus transcripts. This process, known as intraparticle genome transcription, occurs in other icosahedral dsRNA viruses, such as reoviruses and totiviruses (28–36). However, in birnaviruses, the surface pore might be obstructed by unresolved surface loops of VP2 at the 5-fold axis (12, 13). Previous studies have described the capability of purified birnavirus particles to perform intraparticle genome synthesis (37, 38). In contrast, it has also been suggested that the interior RNPs could be released from the capsid during the endocytosis and synthesize viral transcripts in cellular virus factories (39–41). Whether the RNP functions as a capsid-independent transcription complex in situ remains a pending question.

Considering the importance and unrevealed multifunctionality of the birnavirus capsid, it is critical to determine its infectious large T=13 particles using cryogenic electron microscopy (cryo-EM) single particle analysis (SPA). To date, only the cryo-EM structure of IBDV has been resolved (12). In this study, we present the first cryo-EM structure of the salmonid IPNV, providing new insights into the structural mechanisms of VP2 that are essential for infection, particle formation, and intraparticle genome synthesis.

## Results and Discussions

### Cryo-EM structure of T=13 IPNV capsid

The first infectious IPNV capsid structure was determined at a resolution of 2.75 Å using cryo-EM single particle analysis with imposed icosahedral symmetry (Supplementary Fig. S1). The achieved resolution enabled the building of an atomic model of the IPNV capsid.

The IPNV capsid employs a T=13 *laevo* icosahedral lattice, consisting of 60 copies of the asymmetric unit (highlighted with red dotted line in Fig. 1A). Each asymmetric unit is composed of 13 VP2 subunits (labeled (a) - (m) in Fig. 1B). These VP2 subunits form trimers, such as the one composed of subunits (a), (b), and (c) (Fig. 1B), which may serve as building blocks of the birnavirus capsid. From another perspective, five subunits (a) form a pentagonal unit at the 5-fold axis, while the remaining subunits contribute to hexagonal units in the capsid (Fig. 1B). The atomic models of the 13 VP2s were built within the asymmetric unit (Fig. 1B) and subsequently superimposed to observe their structural diversity (Fig. 1C). The VP2 subunits are organized into three domains: the α-helix-rich N and C-terminal base (B) domain, located on the capsid interior; the conserved β-barrel jellyroll fold shell (S) domain; and the unique β-barrel jellyroll fold projection (P) domain, which is situated on the capsid exterior (Fig. 1C). The P domains of the trimer unit form unique protrusions on the capsid surface. While the overall structure of the VP2 subunits is similar, notable variations are observed in the C-terminal region of the B domain (residues 429-435) and the variable loop region of the S domain (residues 110-117) (Fig. 1C).

**Fig. 1.**
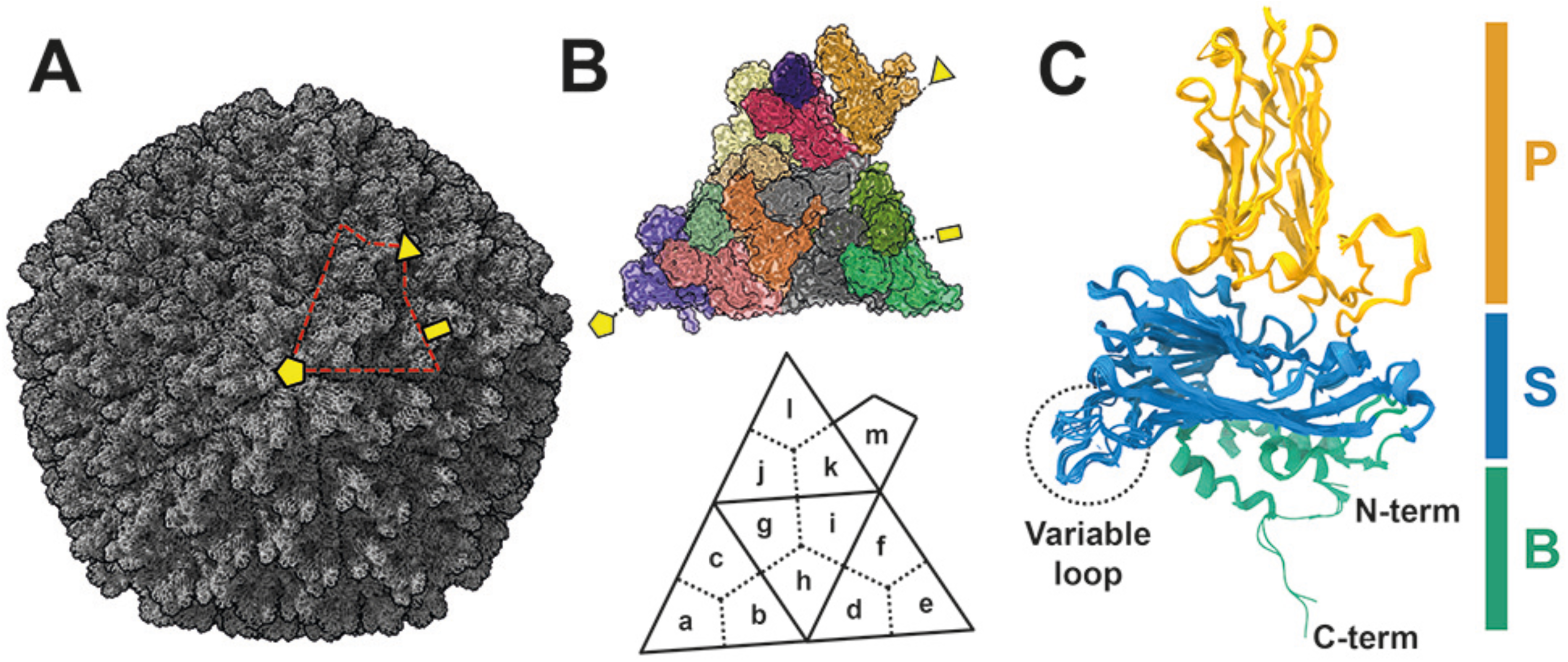
Capsid structure of T=13 infectious IPNV. A) Overall cryo-EM structure of IPNV. The asymmetric unit is encircled by a red dotted line. B) Subunits (a) - (m) within the T=13 asymmetric unit, each depicted in a different color. C) Atomic structure of the VP2 subunits, with all VP2 subunits superimposed. The P, S, and B domains are colored orange, light blue, and light green, respectively. Variable loops are shown in a black dotted circle. The 2-, 3-, and 5-fold axes are shown by a yellow rectangle, triangle, and pentagon in panels (A) and (B).

### Structural comparison between IPNV, IBDV, and other related viruses

The conserved jellyroll structure of the birnavirus VP2 S domain suggests an evolutionary relationship with ssRNA tetraviruses/nodaviruses within the order *Nodamuvirales* (13). Indeed, the jellyroll fold in VP2 of IPNV is similar to those found in capsid proteins of viruses from the *Nodamuvirales* order and *Tetraviridae* family, including Lake Sinai virus, Nudaurelia capensis omega virus, and Pariacoto virus (Supplementary Table S1, Dali score > 10). These viruses also display birnavirus-like P-domains on their capsid surfaces (42–46), implying the importance of the P-domain in host-specific receptor-binding across a broad range of icosahedral jellyroll viruses. The surface loops of the P-domain are structurally different between the T=1 subviral particles of IPNV and IBDV, demonstrating their involvement in host-specific infection (16). A comparison between the cryo-EM structures of the infectious T=13 IBDV and IPNV particles similarly reveals differences in these surface loops (Figs. 2A-B), reinforcing earlier observations made from the T=1 structures. Both IPNV and IBDV form four corresponding surface loops: L1 (IPNV: 206-222, IBDV: 220-224), L2 (IPNV: 206-222, IBDV: 220-224), L3 (IPNV: 277-287, IBDV: 278-287), and L4 (IPNV: 313-323, IBDV: 317-323), while an additional loop, L4’ (326-333) is unique to IPNV (Figs. 2C-D). These surface loops represent the most structurally divergent regions, especially the IPNV L4’ region (Figs. 2B-D). These updated cryo-EM structures of the infectious IPNV and IBDV surfaces reveal more accurate and native insights into the structural differences that may influence host tropism and virulence.

**Fig. 2.**
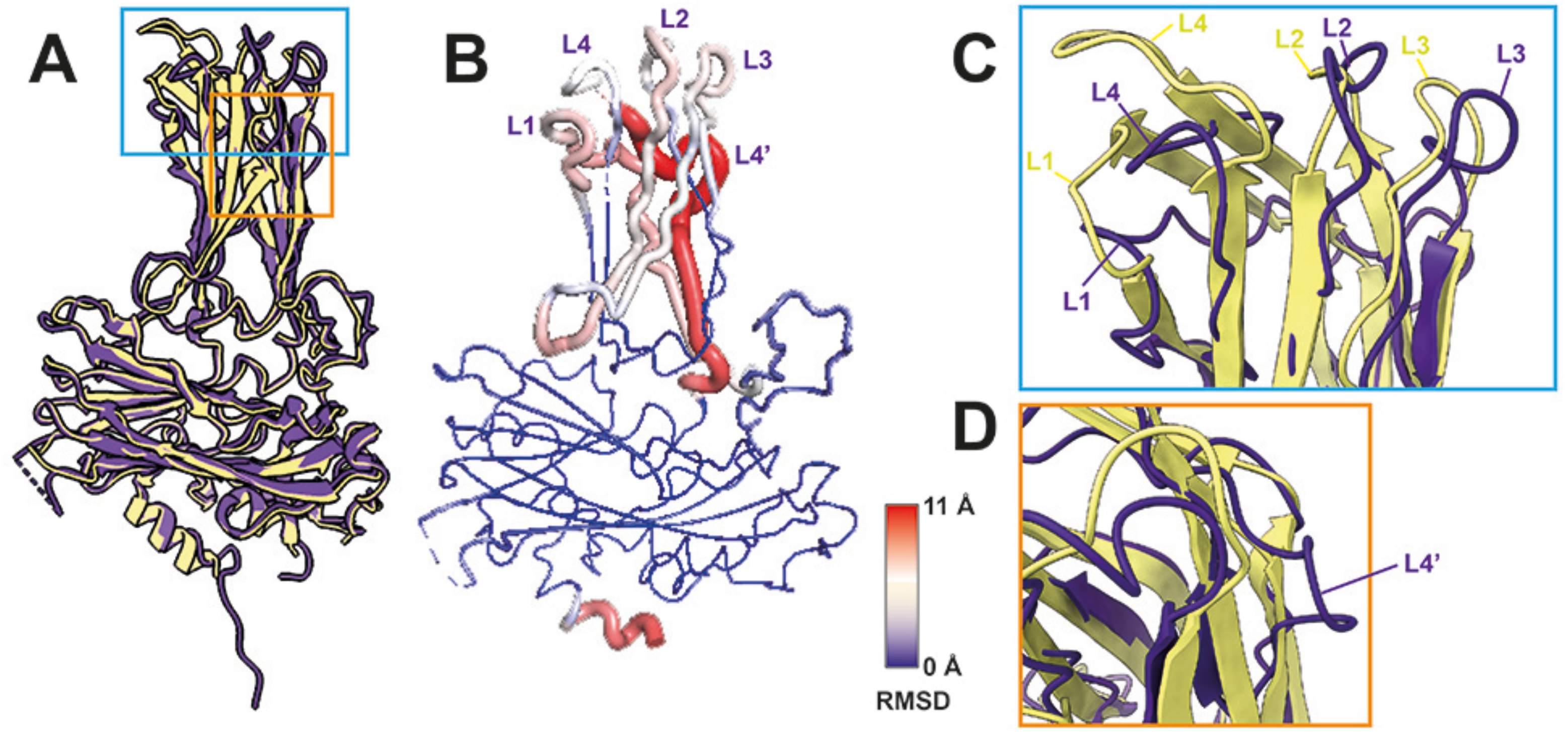
Structural comparisons of VP2 in T=13 IPNV and IBDV particles. A) Superimposed VP2 structures (subunit (a)) of IPNV (purple) and IBDV (yellow). B) Residual RMSD values between IPNV and IBDV VP2 structures. C, D) Close-up views of the aligned VP2 P-domains of IPNV (purple) and IBDV (yellow). Five IPNV (L1-4, L4’) and four IBDV (L1-4) surface loops are labeled.

### Unique structural interface of infectious T=13 IPNV particle

Structural comparison between T=1 and T=13 IPNV particles reveal differences in the variable loop regions of VP2 (Fig. 3A), which also exhibit structural diversity among the 13 VP2 subunits in the T=13 capsid (Fig. 1C). The structure of the VP2 variable loop in the subviral particle was not fully resolved (16), similar to the variable loop of the VP2 subunit (a) that is closest to the 5-fold axis in the T=13 particle (Fig. 3A). However, the variable loop of the other 12 VP2 subunits in the asymmetric unit adopt either an up or down conformation (Fig. 3B), contributing to the formation of the necessary structural interfaces within the hexagonal capsomers in the T=13 capsid lattice (Fig. 3C). Six variable loops of the VP2 subunits are located at the center of each hexagonal capsomer, with three up and three down conformations (Fig. 3B). The Tyr115 residues of the three up conformations largely contribute to forming the interface (Fig. 3C-D). A similar structural interface, mediated by three Tyr118 residues in the up and down conformations of the variable loops (12), is observed in the hexagonal capsomer of IBDV, highlighting their critical role in forming the T=13 infectious particle in birnaviruses. The structural variations observed in the variable loops in the T=13 IPNV capsid arise from the formation of necessary hexagonal capsomers, which are absent in the T=1 subviral particle. Further, these observations align with the structural resemblance between the VP2 subunit (a), which forms a pentagonal unit at the 5-fold axis, and the VP2 subunit in the T=1 subviral particle, which consists solely of pentagonal units (Fig. 3A).

**Fig. 3.**
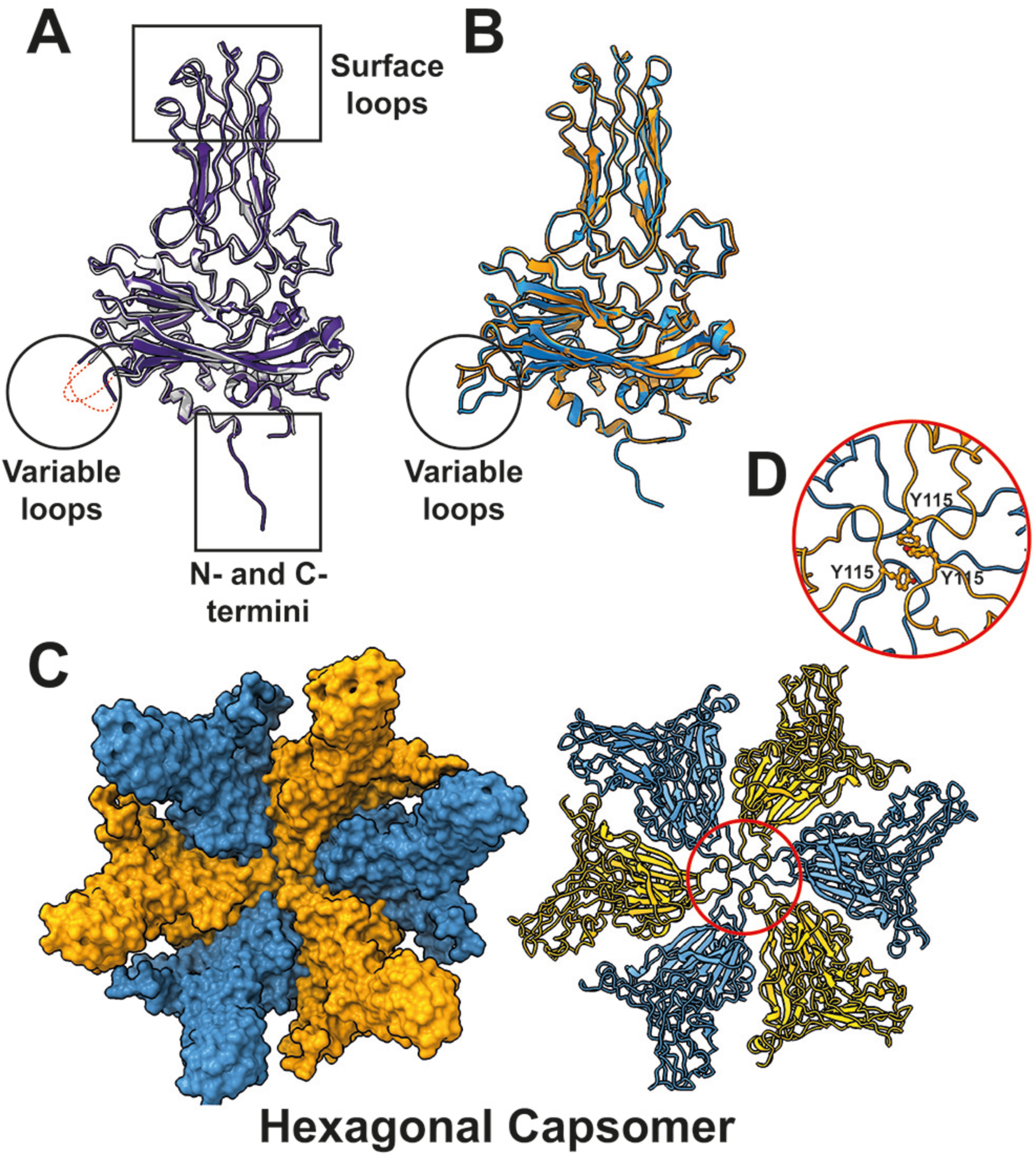
Structure and unique subunit interface of the hexagonal capsomer in T=13 IPNV capsid. A) Structural comparison of VP2 subunits between the T=1 subviral particle (gray, PDBID: 3IDE) and the T=13 infectious particle (purple, subunit (a)) of the IPNV capsid. Large parts of the VP2 variable loop (red dotted lines) are unresolved in these VP2 structures. Structural variations are observed in their variable loops (black circle) and N- and C-termini (black rectangle). Minor structural variations are also observed in surface loops (black rectangle). B) Comparison of the “up” and “down” conformations of the VP2 variable loops in the T=13 infectious particle. The VP2 subunit with the “down” conformation is shown in light blue, while the “up” conformation is orange. C) Structure of the hexagonal capsomer, where six VP2 subunits form hexagonal capsomers in the T=13 capsid, displaying the “down” and “up” conformations of the variable loops. D) Close-up view of the variable loops at the center of the hexagonal capsomer. The Y115 residues in the three VP2 subunits with the “up” conformation contribute to forming the quasi-six fold in the hexagonal capsomer.

### C-terminal interlocking and possible assembly intermediates

The IPNV atomic model, based on residues Thr7 to Asn428 from the T=1 subviral particle crystal structure (Fig. 3A) (16), revealed an extra intensity corresponding to the VP2 C-terminus in some VP2 subunits of the reconstructed T=13 IPNV model. Specifically, this intensity was observed from Glu429 to Ser435 in subunit (a), Glu429 to Ile430 in subunit (b), and Glu429 to Ser434 in subunit (c) (Supplementary Fig. S2). The other VP2 subunits (d) - (m) were modeled up to Asn428, consistent with the T=1 subviral particle. Because of this, the C-terminal extension in the 13 VP2 subunits is structurally variable (Fig. 1B). This extension was previously unresolved in the T=13 IBDV infectious particle and T=1 IPNV subviral particle (12, 16), but its existence has been linked to a possible interlocking role between neighboring VP2 3-fold trimer units in the T=1 IBDV subviral particle (19). In the T=13 IPNV infectious particle, the C-terminal extensions of subunits (a) and (c) interlock with neighboring subunits on the capsid interior (Fig. 4A). The C-terminal region of subunit (a) (residues Thr431, Phe433 - Ser435) interlocks with the adjacent subunit (a)’ at the N-terminal region (residue Lys5), at one α-helix of the B domain (residue Tyr398) and at the C-terminal region (residue Val426) (Fig. 4B, Supplementary Fig. S2A). In contrast, the C-terminal region of subunit (c) (residues Thr431, Phe433) interlocks with the C-terminal region (residues Val426, Glu429) of the adjacent subunit (b) (Fig. 4C, Supplementary Fig. S2B). The shorter C-terminal extension of subunit (b) however, does not orient to the neighboring subunit. Instead, it is located close to the C-terminal extension of subunit (c) (light blue in Figs. 4C, Supplementary Fig. S2B), thus contributing to the interlocking nevertheless. As a result of these interlocking interactions, the five VP2 trimers form a penton protein-protein network complex (Fig. 4A). Long terminal extensions and similar interlocking mechanisms have also been identified in larger icosahedral dsRNA viruses, such as toti-like viruses, picobirnaviruses, megabirnavirus, quadrivirus, and reoviruses that have longer non-segmented or segmented genomes (28, 32, 47–53), and *Totiviridae* viruses that have shorter non-segmented genomes (30, 31, 54). These interlocking networks are believed to stabilize the virus capsid against the internal pressure from their viral genomes. This suggests that the C-terminal extensions of the relatively large and bi-segmented IPNV may have a similar function in stabilizing its capsid.

**Fig. 4.**
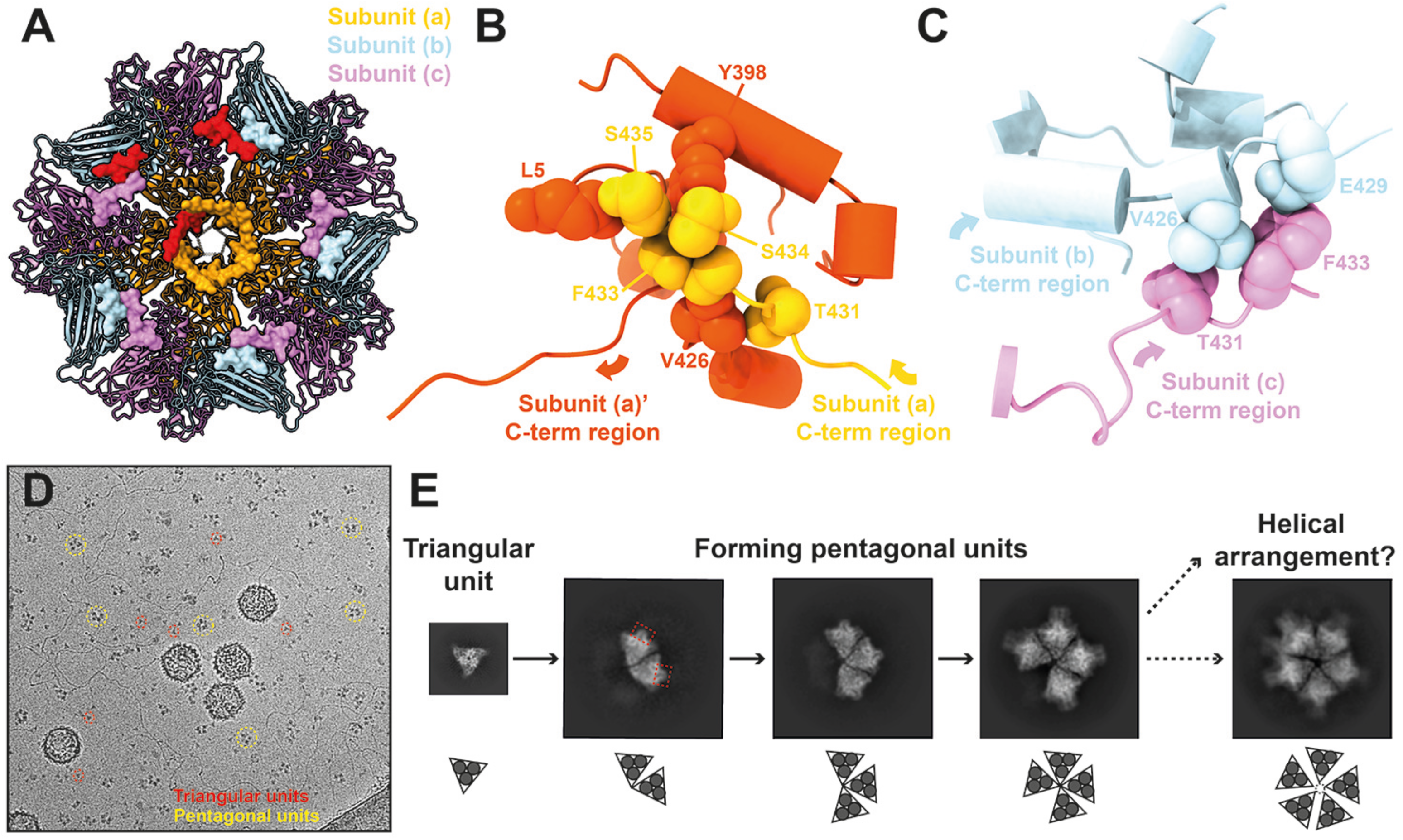
C-terminal extension of VP2 subunits and the intrasubunit interlocking in pentagonal structural units. A) C-terminal interlocking network of VP2 subunits at the interior of a 5-fold vertex. Five VP2 subunits from each of the (a), (b), and (c) subunits form a pentagonal structural unit at the 5-fold vertex of the T=13 IPNV capsid. The C-terminal extensions of these subunits are shown in surface representation. Representative C-terminal extensions from subunits (a), (b) and (c) are shown in red. B) Close-up view of the C-terminal extension region between adjacent subunits (a) and (a)’. C) Close-up view of the C-terminal extension region between adjacent subunits (b) and (c). Interacting amino acid residues are labeled in panels B and C. D) Triangular and pentagonal units of IPNV in a raw cryo-EM micrograph. E) 2D class-averages of the triangular and pentagonal units. Red dotted rectangles show protruded features of the pentagonal units.

The C-terminal extension could also be important for assembling the T=13 capsid. During the maturation process of the IPNV polyprotein, the C-terminal region of pVP2 is expected to be cleaved in multiple sites (Ala442, Ala486, and Ala495) to form mature VP2 (8). Although the exact length of the mature VP2 is not fully clear, it is anticipated that the C-terminus should exist at the Ala442 residue in the mature IPNV particle. Recombinant expression of only mature VP2 (residues 1-422) results in the formation of a T=1 subviral particle, whereas the inclusion of the C-terminal region of pVP2 (residues 1-466) and the C-terminal domain of VP3 results in the formation of T=13 and T=1 icosahedrons as well as tubular structures (14), highlighting the importance of the previously unresolved C-terminal extension of VP2 (residues 429-435) (Fig. 4). The structure and the interlocking function of the C-terminal extension could be critical for assembling the T=13 capsid and for forming the required intermediate building blocks. The observed interlocked penton unit in the IPNV T=13 capsid does not exist in the T=1 subviral particle. The C-terminal interlocking could prevent the incorrect formation of T=1 capsids by assembling penton structural units during pVP2 maturation and assembly.

Many triangular and partial or complete pentagonal units were observed in the raw cryo-EM images of the IPNV sample (Fig. 4D). The 2D-class averaging of these units revealed that the triangular units correspond to individual VP2 trimers, while the partial and complete pentagonal units consist of 2-5 trimer units (Fig. 4E). The formation of these pentagonal units from five triangular units aligns with previously speculated models for IBDV assembly (15, 55). Notably, in the 2D-classes of 2-4 trimers, the central region of the complex remains relatively closed, whereas the complete penton unit exhibits a more open structure (Fig. 4E). Comparison of the 5-fold pore structures composed of five VP2 subunits in the T=1 and T=13 IPNV capsids – excluding the variable loop region (residues 110-117) – reveals a larger 5-fold pore size in the T=13 penton unit (19.8 Å in radius) compared to the T=1 penton unit (18.9 Å in radius) (Supplementary Fig. S3). This suggests that the penton unit in the T=13 is more sparsely packed, potentially contributing to the observed larger pore in the full pentagonal unit (Fig. 4E). We also hypothesize that the helical structure of the VP2 trimers may form preferentially if the pore does not expand during pentagonal unit assembly, possibly due to spatial constraints for accommodating the fifth VP2 trimer. Although the 3D reconstructions of these triangle and pentagonal units are performed, the clear 3D model has not been built yet because of preferred orientation. Additional protruding features are visible in the trimer units in the 2D projections of the pentagonal units, which are absent in the triangular units (red dotted rectangles in Fig. 4E). One speculation is that these protrusions may represent part of the C-terminal regions of pVP2 subunits that need to be rearranged during the transition from triangular to pentagonal units. Further studies on birnavirus assembly intermediates are essential to elucidate the role of the C-terminal extensions and the formation of pentagonal units.

### Potential gate structure of IPNV

Five VP2 subunits (subunits (a) in Fig. 1B) form a 5-fold pore structure in the IPNV capsid. As discussed in previous studies of the T=13 IBDV capsid (12, 13), this pore may be obstructed by unresolved variable loops from the five VP2 subunits at each 5-fold axis. As aforementioned, the T=13 IPNV capsid also shows 5-fold pore structures with unresolved variable loops likely positioned within the pore (Fig. 4A, Supplementary Fig. S3). The 5-fold focused asymmetric reconstruction of the IPNV pore indicates more pronounced obstruction, though a clear structure is not discernible, most likely due to the different conformations of the variable loops (Fig. 5A). Despite ongoing debate about intraparticle genome synthesis in Birnaviruses, the IPNV pore shares similar characteristics with other dsRNA viruses known to facilitate intraparticle genome synthesis. The VP2 subunit exhibits two positively charged residues, Lys159 and Arg425, on the pore surface, which form clusters of positively charged regions (Figs. 5B-C). Similar clusters in other icosahedral dsRNA viruses are believed to be important for recruiting negatively charged NTPs to facilitate RdRp-mediated intraparticle genome synthesis (28, 30–32, 47). Moreover, in single-layered dsRNA viruses that infect multicellular hosts (metazoa), including artiviruses (arthropod totiviruses), mammalian picornaviruses, and fungal megabirnavirus, the gate of their pore is either closed or obstructed by flexible loops or additional proteins (32, 47, 48, 51, 53). Conclusively, these structural features of the salmonid IPNV capsid pore are akin to the previously observed pore structure of the metazoan dsRNA viruses. Understanding the genome synthesis mechanisms in these viruses necessitates determining the interior RNP structures (VP1 and VP3). Although asymmetric reconstruction methods were attempted according to previous dsRNA virus structural studies, a clear 3D model of the genome and RNP structures of IPNV could not be determined, most likely due to heterogeneity. However, 2D class-averaged particle images of the capsid-subtracted dataset reveal a non-typical organization of IPNV RNPs, differing from the partially spooled dsRNA genomes seen in *Totivirida*e and *Reoviridae* viruses (Fig. 5D) (28, 29, 34, 35, 56). This implies that IPNV employs a different genome packaging mechanism for the intraparticle genome synthesis.

**Fig. 5.**
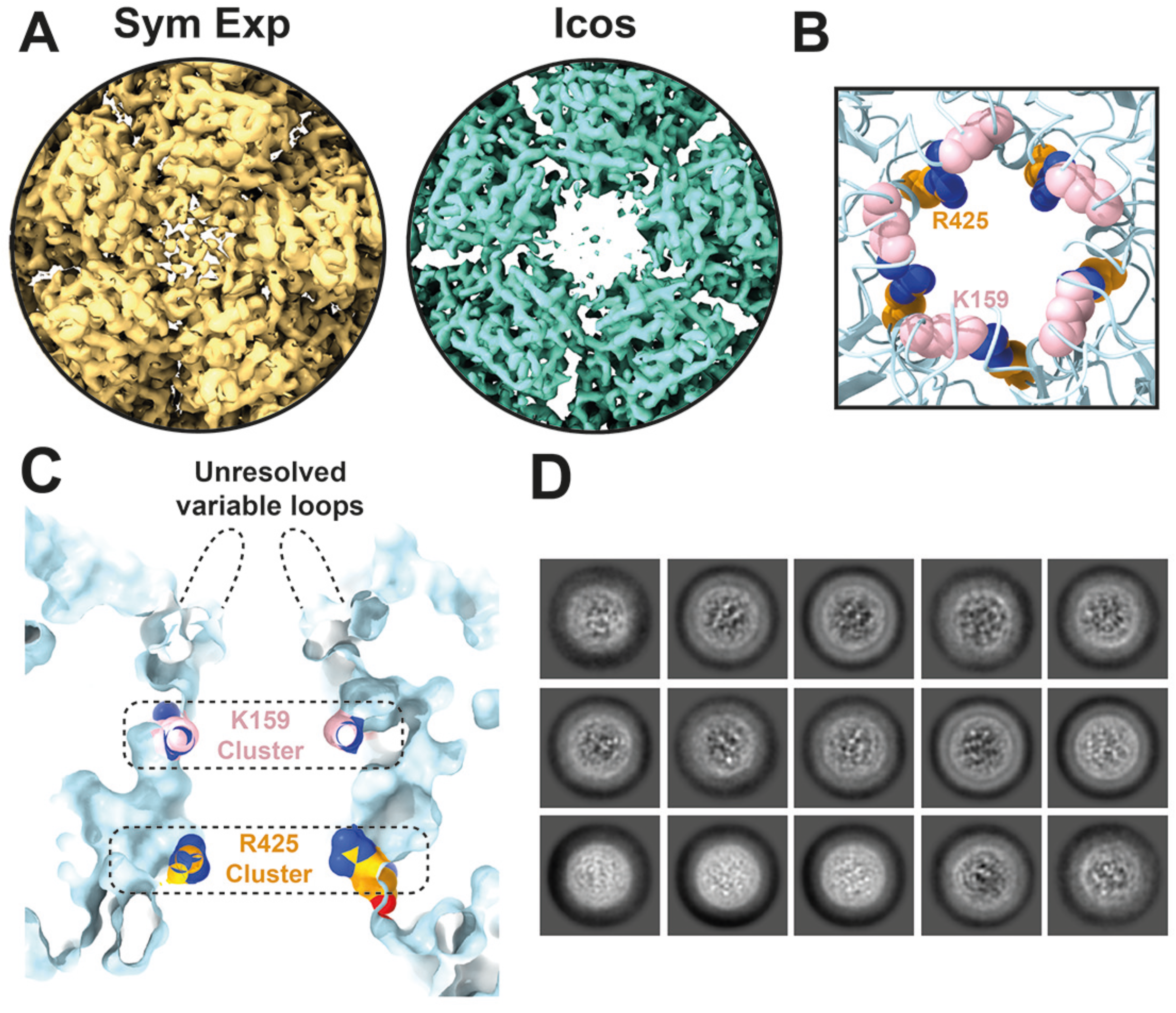
Potential gate for intraparticle genome transcription in Birnavirus. A) 5-fold pore structure as revealed by icosahedrally averaged (Icos) and symmetry expanded (Sym Exp) cryo-EM reconstructions of the IPNV capsid. The contour level of each cryo-EM map is adjusted to 5 sigma. B) Top view of the 5-fold pore. B) Cross-section of the 5-fold pore. B, C) Residues 110-117 (unresolved variable loops) of the VP2 are removed for visualization. Positively charged residues of the gate, five K159 and R425 residues, are colored pink and orange. D) Representative particle images obtained from capsid-subtracted 2D-class averaging.

## Materials and Methods

### Cell culture and IPNV sample preparation

The IPNV strain (IPNV-L5) used in this study was isolated from head kidney samples obtained from cultured fish originating from sites with suspected IPNV. The initial isolation was performed in CHSE-214 cells and the seed virus stock used for propagating IPNV-L5 samples in this study was propagated three times in RTG2 cells. The VP2 gene of IPNV L5 varied from the classical virulent isolate of IPNV (Genbank ID: AY379740) at 12 positions, listed in Supplementary Table S2.

The purification protocol for IPNV particles was optimized from previously reported papers (57, 58). Infected culture fluid (ICF) was centrifuged at 5,000 x*g*, 4°C for 30 min to remove cell debris. Solid PEG 20,000 and sodium chloride were added to the ICF to achieve final concentrations of 5% (w/v) and 2.2% (w/v), respectively, and the mixture was stirred overnight at 4°C. The virus-containing pellet was obtained by centrifuging the ICF at 15,300 x*g*, 4°C for 60 min. The pellet was resuspended in 1 mL of PBS (-). The sample was applied on a 15 - 45% (w/v) sucrose gradient and ultracentrifuged at 130,000 x*g*, 4°C for 2.5 hr. The SDS-PAGE result of the sucrose gradient fractions is shown in Supplementary Fig. S1A. The fractions containing filled particles with VP1, VP2, and VP3 protein bands were pooled and diluted in PBS (-) up to 13 mL. The diluted virus fractions were ultracentrifuged at 130,000 x*g*, 4°C for 2 hr. The virus pellet was resuspended in 50 µL of PBS (-), and then concentrated to 20 µL. The final protein concentration of the sample was calculated to be 1.08 mg/mL, which was used for cryo-EM grid preparation.

### Cryo-EM data acquisition and map reconstruction

The data acquisition parameters are listed in Supplementary Table S3. Cryo-EM grids were obtained by flash-freezing 3 µL of purified IPNV particles on glow-discharged holey carbon grids (Quantifoil R2/2, Cu 300 mesh; Quantifoil Micro Tools GmbH). The grids were plunge-frozen in liquid ethane after blotting for 3 s under 4°C, 100% humidity using a Mark IV Vitrobot (Thermo Fisher Scientific). The frozen grids were initially screened with a Glacios cryo-EM (Thermo Fisher Scientific) at the Uppsala cryo-EM center. The complete dataset was collected with a Titan Krios (Thermo Fisher Scientific) installed with a Gatan K3 BioQuantum detector and an electron energy filter (20 eV slit width) at the SciLifeLab Cryo-EM facility. Image movies (30 frames per movie) were acquired in counted superresolution mode at a nominal magnification of 81,000x, corresponding to a pixel size of 1.058 Å/pixel. The total electron exposure was 26.8 e^-^/Å^2^ over 1.34 s per movie. Defocus values were randomly selected in 0.2 µm increments within the range of 0.7 to 1.5 µm underfocus for each movie.

A total of 27,945 movies were collected for three-dimensional (3D) cryo-EM model reconstruction. Image processing and 3D reconstruction were conducted using CryoSPARC version 4.3.1 (59) and a local CPU/GPU computer cluster. Initial preprocessing involved patch motion correction using frames 3-27, followed by contrast transfer function (CTF) estimation via patch CTF. Template picking and 2D classifications selected 35,498 particles, including 12,898 dsRNA-genome-filled particles. The 3D reconstructions were performed on downsampled particles (1.284 Å/pixel). An initial model was generated from 6,471 particles via ab initio reconstruction. The best achievable resolution was obtained by processing all selected 35,498 particles using the Homogeneous Refinement option with imposed icosahedral symmetry. During refinement, per-particle defocus and per-group CTF parameters were optimized, with spherical aberration, beam tetrafoil, and anisotropic magnification fitting enabled. Ewald sphere correction was also performed. The final 3D map of IPNV L5 achieved a resolution of 2.75 Å based on Fourier shell correlation (FSC) at a 0.143 cutoff (60) (Supplementary Fig. S1C). Due to the large size of the particle, 70 nm in diameter, the block-based reconstruction was also performed (12, 61, 62); however, no significant improvement was observed. The obtained cryo-EM map was used for building atomic models of the VP2 subunits. The 2D classification of the IPNV interior structure was performed using capsid-subtracted images of the 12,898 filled particles. Asymmetric reconstruction was done using a previously described symmetry expansion and local reconstruction method for icosahedral dsRNA viruses (28, 56). A mask covering one 5-fold vertex was applied during local reconstruction.

### Atomic modeling, refinement, and rendering of IPNV capsid

The initial atomic model of the VP2 CP of IPNV-L5 was predicted from the VP2 sequence using AlphaFold2 (63). The highest-scoring predicted model was then first fitted into the subunit (a) region (the closest subunit to the 5-fold vertex) of the cryo-EM map using UCSF Chimera (64). The atomic model was then manually refined using Coot version 1.0.06 (65) and further automatically refined using PHENIX 1.20.1 (66) in iterative cycles. The refined atomic model of VP2 was then manually fitted to each of the other 12 VP2 subunit locations (subunits (b) - (m)) within an asymmetric unit of the icosahedral map. These 12 VP2 were also refined using Coot and Phenix software. Validation statistics of the atomic models are shown in Supplementary Table S3. For rendering the cryo-EM maps and the atomic models, the UCSF Chimera and ChimeraX were used (64, 67).

### Structural comparison

The resolved structure of IPNV VP2 and corresponding CPs from other viruses that employ a jellyroll fold were extensively compared using the DALI server (68). The structural alignment of VP2 subunits from IBDV and IPNV capsid structures was generated using the TM-align algorithm (69) and the residual RMSD was calculated in PyMOL as described previously (26, 28, 69).

## Data availability

The cryo-EM map of the IPNV-L5 is available in the EMDB database, entry EMD-51321. The atomic models of the CPs are available in the PDB database, entry 9GG2.

## Acknowledgments

The data was collected at the Cryo-EM Swedish National Facility funded by the Knut and Alice Wallenberg, Erling Persson Family, and Kempe Foundations, SciLifeLab, Stockholm University and Umeå University. We thank Dustin Morado for help with data acquisition, and Jakob Klefenberg for helping with data analysis. Funding was provided by the following agencies: Research Council of Norway (to Ø.E., grant number: 324266), the Swedish Research Council (to K.O., grant number: 2018-03387 and 2023-01857; to A.M., grant number: 2022-00236), the Swedish Foundation for International Cooperation in Research and Higher Education (STINT) (to K.O., grant number: JA2014-5721), FORMAS research grant from the Swedish Research Council for Environment, Agricultural Sciences and Spatial Planning (to K.O., grant number: 2018-00421 and 2022-02347), Royal Swedish Academy of Sciences (to K.O., grant number: BS2018-0053). The funders had no role in study design, data collection and interpretation, or the decision to submit the work for publication.

## References

Yang Z, He B, Lu Z, Mi S, Jiang J, Liu Z, Tu C, Gong W. 2021. Mammalian birnaviruses identified in pigs infected by classical swine fever virus. Virus Evol 7:veab084.

Evensen O, Rimstad E. 1990. Immunohistochemical identification of infectious pancreatic necrosis virus in paraffin-embedded tissues of Atlantic salmon (Salmo salar). J Vet Diagn Invest 2:288–293.

Wolf K, Snieszko SF, Dunbar CE, Pyle E. 1960. Virus nature of infectious pancreatic necrosis in trout. Proc Soc Exp Biol Med 104:105–108.

Evensen Ø, Leong J-AC. 2013. DNA vaccines against viral diseases of farmed fish. Fish Shellfish Immunol 35:1751–1758.

Giambrone JJ, Donahoe JP, Dawe DL, Eidson CS. 1977. Specific suppression of the bursa-dependent immune system of chicks with infectious bursal disease virus. Am J Vet Res 38:581–583.

Mahgoub HA, Bailey M, Kaiser P. 2012. An overview of infectious bursal disease. Arch Virol 157:2047–2057.

Nobiron I, Galloux M, Henry C, Torhy C, Boudinot P, Lejal N, Da Costa B, Delmas B. 2008. Genome and polypeptides characterization of Tellina virus 1 reveals a fifth genetic cluster in the Birnaviridae family. Virology 371:350–361.

Da Costa B, Soignier S, Chevalier C, Henry C, Thory C, Huet J-C, Delmas B. 2003. Blotched snakehead virus is a new aquatic birnavirus that is slightly more related to avibirnavirus than to aquabirnavirus. J Virol 77:719–725.

Da Costa B, Chevalier C, Henry C, Huet J-C, Petit S, Lepault J, Boot H, Delmas B. 2002. The capsid of infectious bursal disease virus contains several small peptides arising from the maturation process of pVP2. J Virol 76:2393–2402.

Mertens J, Casado S, Mata CP, Hernando-Pérez M, de Pablo PJ, Carrascosa JL, Castón JR. 2015. A protein with simultaneous capsid scaffolding and dsRNA-binding activities enhances the birnavirus capsid mechanical stability. Sci Rep 5:13486.

Bahar MW, Sarin LP, Graham SC, Pang J, Bamford DH, Stuart DI, Grimes JM. 2013. Structure of a VP1-VP3 complex suggests how birnaviruses package the VP1 polymerase. J Virol 87:3229–3236.

Bao K, Qi X, Li Y, Gong M, Wang X, Zhu P. 2022. Cryo-EM structures of infectious bursal disease viruses with different virulences provide insights into their assembly and invasion. Sci Bull (Beijing) 67:646–654.

Coulibaly F, Chevalier C, Gutsche I, Pous J, Navaza J, Bressanelli S, Delmas B, Rey FA. 2005. The birnavirus crystal structure reveals structural relationships among icosahedral viruses. Cell 120:761–772.

Saugar I, Irigoyen N, Luque D, Carrascosa JL, Rodríguez JF, Castón JR. 2010. Electrostatic interactions between capsid and scaffolding proteins mediate the structural polymorphism of a double-stranded RNA virus. J Biol Chem 285:3643– 3650.

Lee C-C, Ko T-P, Chou C-C, Yoshimura M, Doong S-R, Wang M-Y, Wang AH-J. 2006. Crystal structure of infectious bursal disease virus VP2 subviral particle at 2.6A resolution: implications in virion assembly and immunogenicity. J Struct Biol 155:74–86.

Coulibaly F, Chevalier C, Delmas B, Rey FA. 2010. Crystal structure of an Aquabirnavirus particle: insights into antigenic diversity and virulence determinism. J Virol 84:1792–1799.

Chevalier C, Lepault J, Erk I, Da Costa B, Delmas B. 2002. The maturation process of pVP2 requires assembly of infectious bursal disease virus capsids. J Virol 76:2384–2392.

Oña A, Luque D, Abaitua F, Maraver A, Castón JR, Rodríguez JF. 2004. The C-terminal domain of the pVP2 precursor is essential for the interaction between VP2 and VP3, the capsid polypeptides of infectious bursal disease virus. Virology 322:135–142.

Garriga D, Querol-Audí J, Abaitua F, Saugar I, Pous J, Verdaguer N, Castón JR, Rodriguez JF. 2006. The 2.6-Angstrom structure of infectious bursal disease virus-derived T=1 particles reveals new stabilizing elements of the virus capsid. J Virol 80:6895–6905.

Munke A, Kimura K, Tomaru Y, Wang H, Yoshida K, Mito S, Hongo Y, Okamoto K. 2022. Primordial Capsid and Spooled ssDNA Genome Structures Unravel Ancestral Events of Eukaryotic Viruses. MBio 13:e0015622.

Santi N, Vakharia VN, Evensen Ø. 2004. Identification of putative motifs involved in the virulence of infectious pancreatic necrosis virus. Virology 322:31–40.

Mutoloki S, Jøssund TB, Ritchie G, Munang’andu HM, Evensen Ø. 2016. Infectious pancreatic necrosis virus causing clinical and subclinical infections in atlantic salmon have different genetic fingerprints. Front Microbiol 7:1393.

Song H, Santi N, Evensen O, Vakharia VN. 2005. Molecular determinants of infectious pancreatic necrosis virus virulence and cell culture adaptation. J Virol 79:10289–10299.

Qi X, Zhang L, Chen Y, Gao L, Wu G, Qin L, Wang Y, Ren X, Gao Y, Gao H, Wang X. 2013. Mutations of residues 249 and 256 in VP2 are involved in the replication and virulence of infectious Bursal disease virus. PLoS ONE 8:e70982.

Zhu L, Wang X, Ren J, Porta C, Wenham H, Ekström J-O, Panjwani A, Knowles NJ, Kotecha A, Siebert CA, Lindberg AM, Fry EE, Rao Z, Tuthill TJ, Stuart DI. 2015. Structure of Ljungan virus provides insight into genome packaging of this picornavirus. Nat Commun 6:8316.

Wang H, Munke A, Li S, Tomaru Y, Okamoto K. 2022. Structural Insights into Common and Host-Specific Receptor-Binding Mechanisms in Algal Picorna-like Viruses. Viruses 14.

Munke A, Kimura K, Tomaru Y, Okamoto K. 2020. Capsid structure of a marine algal virus of the order picornavirales. J Virol 94.

Wang H, Marucci G, Munke A, Hassan MM, Lalle M, Okamoto K. 2024. High-resolution comparative atomic structures of two Giardiavirus prototypes infecting G. duodenalis parasite. PLoS Pathog 20:e1012140.

Bao K, Zhang X, Li D, Sun W, Sun Z, Wang J, Zhu P. 2022. In situ structures of polymerase complex of mammalian reovirus illuminate RdRp activation and transcription regulation. Proc Natl Acad Sci USA 119:e2203054119.

Grybchuk D, Procházková M, Füzik T, Konovalovas A, Serva S, Yurchenko V, Plevka P. 2022. Structures of L-BC virus and its open particle provide insight into Totivirus capsid assembly. Commun Biol 5:847.

Stevens A, Muratore K, Cui Y, Johnson PJ, Zhou ZH. 2021. Atomic Structure of the Trichomonas vaginalis Double-Stranded RNA Virus 2. MBio 12.

Okamoto K, Ferreira RJ, Larsson DSD, Maia FRNC, Isawa H, Sawabe K, Murata K, Hajdu J, Iwasaki K, Kasson PM, Miyazaki N. 2020. Acquired Functional Capsid Structures in Metazoan Totivirus-like dsRNA Virus. Structure 28:888–896.e3.

He Y, Shivakoti S, Ding K, Cui Y, Roy P, Zhou ZH. 2019. In situ structures of RNA-dependent RNA polymerase inside bluetongue virus before and after uncoating. Proc Natl Acad Sci USA 116:16535–16540.

Wang X, Zhang F, Su R, Li X, Chen W, Chen Q, Yang T, Wang J, Liu H, Fang Q, Cheng L. 2018. Structure of RNA polymerase complex and genome within a dsRNA virus provides insights into the mechanisms of transcription and assembly. Proc Natl Acad Sci USA 115:7344–7349.

Zhang X, Ding K, Yu X, Chang W, Sun J, Zhou ZH. 2015. In situ structures of the segmented genome and RNA polymerase complex inside a dsRNA virus. Nature 527:531–534.

Guglielmi KM, McDonald SM, Patton JT. 2010. Mechanism of intraparticle synthesis of the rotavirus double-stranded RNA genome. J Biol Chem 285:18123–18128.

Spies U, Müller H, Becht H. 1987. Properties of RNA polymerase activity associated with infectious bursal disease virus and characterization of its reaction products. Virus Res 8:127–140.

Mertens PP, Jamieson PB, Dobos P. 1982. In vitro RNA synthesis by infectious pancreatic necrosis virus-associated RNA polymerase. J Gen Virol 59:47–56.

Delgui LR, Rodríguez JF, Colombo MI. 2013. The endosomal pathway and the Golgi complex are involved in the infectious bursal disease virus life cycle. J Virol 87:8993–9007.

Dalton RM, Rodríguez JF. 2014. Rescue of infectious birnavirus from recombinant ribonucleoprotein complexes. PLoS ONE 9:e87790.

Brodrick AJ, Broadbent AJ. 2023. The formation and function of birnaviridae virus factories. Int J Mol Sci 24.

Banerjee M, Speir JA, Kwan MH, Huang R, Aryanpur PP, Bothner B, Johnson JE. 2010. Structure and function of a genetically engineered mimic of a nonenveloped virus entry intermediate. J Virol 84:4737–4746.

Chen N-C, Wang C-H, Yoshimura M, Yeh Y-Q, Guan H-H, Chuankhayan P, Lin C-C, Lin P-J, Huang Y-C, Wakatsuki S, Ho M-C, Chen C-J. 2023. Structures of honeybee-infecting Lake Sinai virus reveal domain functions and capsid assembly with dynamic motions. Nat Commun 14:545.

Helgstrand C, Munshi S, Johnson JE, Liljas L. 2004. The refined structure of Nudaurelia capensis omega virus reveals control elements for a T = 4 capsid maturation. Virology 318:192–203.

Tang L, Johnson KN, Ball LA, Lin T, Yeager M, Johnson JE. 2001. The structure of pariacoto virus reveals a dodecahedral cage of duplex RNA. Nat Struct Biol 8:77–83.

Speir JA, Taylor DJ, Natarajan P, Pringle FM, Ball LA, Johnson JE. 2010. Evolution in action: N and C termini of subunits in related T = 4 viruses exchange roles as molecular switches. Structure 18:700–709.

Wang H, Salaipeth L, Miyazaki N, Suzuki N, Okamoto K. 2023. Capsid structure of a fungal dsRNA megabirnavirus reveals its previously unidentified surface architecture. PLoS Pathog 19:e1011162.

Shao Q, Jia X, Gao Y, Liu Z, Zhang H, Tan Q, Zhang X, Zhou H, Li Y, Wu D, Zhang Q. 2021. Cryo-EM reveals a previously unrecognized structural protein of a dsRNA virus implicated in its extracellular transmission. PLoS Pathog 17:e1009396.

Mata CP, Luque D, Gómez-Blanco J, Rodríguez JM, González JM, Suzuki N, Ghabrial SA, Carrascosa JL, Trus BL, Castón JR. 2017. Acquisition of functions on the outer capsid surface during evolution of double-stranded RNA fungal viruses. PLoS Pathog 13:e1006755.

Okamoto K, Miyazaki N, Larsson DSD, Kobayashi D, Svenda M, Mühlig K, Maia FRNC, Gunn LH, Isawa H, Kobayashi M, Sawabe K, Murata K, Hajdu J. 2016. The infectious particle of insect-borne totivirus-like Omono River virus has raised ridges and lacks fibre complexes. Sci Rep 6:33170.

Duquerroy S, Da Costa B, Henry C, Vigouroux A, Libersou S, Lepault J, Navaza J, Delmas B, Rey FA. 2009. The picobirnavirus crystal structure provides functional insights into virion assembly and cell entry. EMBO J 28:1655–1665.

Reinisch KM, Nibert ML, Harrison SC. 2000. Structure of the reovirus core at 3.6 A resolution. Nature 404:960–967.

Ortega-Esteban Á, Mata CP, Rodríguez-Espinosa MJ, Luque D, Irigoyen N, Rodríguez JM, de Pablo PJ, Castón JR. 2020. Cryo-electron Microscopy Structure, Assembly, and Mechanics Show Morphogenesis and Evolution of Human Picobirnavirus. J Virol 94.

Naitow H, Tang J, Canady M, Wickner RB, Johnson JE. 2002. L-A virus at 3.4 A resolution reveals particle architecture and mRNA decapping mechanism. Nat Struct Biol 9:725–728.

Luque D, Saugar I, Rodríguez JF, Verdaguer N, Garriga D, Martín CS, Velázquez-Muriel JA, Trus BL, Carrascosa JL, Castón JR. 2007. Infectious bursal disease virus capsid assembly and maturation by structural rearrangements of a transient molecular switch. J Virol 81:6869–6878.

Pan M, Alvarez-Cabrera AL, Kang JS, Wang L, Fan C, Zhou ZH. 2021. Asymmetric reconstruction of mammalian reovirus reveals interactions among RNA, transcriptional factor µ2 and capsid proteins. Nat Commun 12:4176.

Carlsson A, Kuznar J, Varga M, Everitt E. 1994. Purification of infectious pancreatic necrosis virus by anion exchange chromatography increases the specific infectivity. J Virol Methods 47:27–35.

Dobos P, Hill BJ, Hallett R, Kells DT, Becht H, Teninges D. 1979. Biophysical and biochemical characterization of five animal viruses with bisegmented double-stranded RNA genomes. J Virol 32:593–605.

Punjani A, Rubinstein JL, Fleet DJ, Brubaker MA. 2017. cryoSPARC: algorithms for rapid unsupervised cryo-EM structure determination. Nat Methods 14:290– 296.

Rosenthal PB, Henderson R. 2003. Optimal determination of particle orientation, absolute hand, and contrast loss in single-particle electron cryomicroscopy. J Mol Biol 333:721–745.

Zhu D, Wang X, Fang Q, Van Etten JL, Rossmann MG, Rao Z, Zhang X. 2018. Pushing the resolution limit by correcting the Ewald sphere effect in single-particle Cryo-EM reconstructions. Nat Commun 9:1552.

Burton-Smith RN, Narayana Reddy HK, Svenda M, Abergel C, Okamoto K, Murata K. 2021. The 4.4 Å structure of the giant Melbournevirus virion belonging to the *Marseilleviridae* family. BioRxiv 10.1101/2021.07.14.452405.

Jumper J, Evans R, Pritzel A, Green T, Figurnov M, Ronneberger O, Tunyasuvunakool K, Bates R, Žídek A, Potapenko A, Bridgland A, Meyer C, Kohl SAA, Ballard AJ, Cowie A, Romera-Paredes B, Nikolov S, Jain R, Adler J, Back T, Hassabis D. 2021. Highly accurate protein structure prediction with AlphaFold. Nature 596:583–589.

Pettersen EF, Goddard TD, Huang CC, Couch GS, Greenblatt DM, Meng EC, Ferrin TE. 2004. UCSF Chimera—a visualization system for exploratory research and analysis. J Comput Chem 25:1605–1612.

Emsley P, Cowtan K. 2004. Coot: model-building tools for molecular graphics. Acta Crystallogr D Biol Crystallogr 60:2126–2132.

Liebschner D, Afonine PV, Baker ML, Bunkóczi G, Chen VB, Croll TI, Hintze B, Hung LW, Jain S, McCoy AJ, Moriarty NW, Oeffner RD, Poon BK, Prisant MG, Read RJ, Richardson JS, Richardson DC, Sammito MD, Sobolev OV, Stockwell DH, Adams PD. 2019. Macromolecular structure determination using X-rays, neutrons and electrons: recent developments in Phenix. Acta Crystallogr D Struct Biol 75:861–877.

Pettersen EF, Goddard TD, Huang CC, Meng EC, Couch GS, Croll TI, Morris JH, Ferrin TE. 2021. UCSF ChimeraX: structure visualization for researchers, educators, and developers. Protein Sci 30:70–82.

Holm L. 2020. DALI and the persistence of protein shape. Protein Sci 29:128– 140.

Zhang Y, Skolnick J. 2005. TM-align: a protein structure alignment algorithm based on the TM-score. Nucleic Acids Res 33:2302–2309.

